# De novo formation of an early endosome during Rab5 to Rab7 transition

**DOI:** 10.1101/2020.07.06.189050

**Authors:** Frode Miltzow Skjeldal, Linda Hofstad Haugen, Duarte Mateus, Dominik Frei, Oddmund Bakke

## Abstract

Rab5 and Rab7a are the main determinants of early and late endosomes and are important regulators of endosomal progression. The transport from early endosomes to late endosome seems to be regulated through an endosomal maturation switch where Rab5 is exchanged with Rab7a on the same endosome. Here we provide new insight into the mechanism of endosomal maturation where we have discovered a stepwise Rab5 detachment, sequentially regulated by Rab7a. The initial detachment of Rab5 is Rab7a independent and demonstrate a diffusion-like exchange between cytosol and endosomal membrane, and the second phase is slower where Rab5 converges into a specific domain that specifically detaches as a Rab5 indigenous endosome. Consequently, we show that early endosomal maturation regulated through the Rab5 to Rab7a switch induce the formation of a new fully functional early endosome. Progression through a stepwise early endosomal maturation regulates the direction of the transport and concomitantly regulates the homeostasis of early endosomes.

## Introduction

Endosomal trafficking is carefully regulated through the mechanisms of endosomal progression. In general, two models have been characterized to explain the trafficking through the endocytic pathway (Helenius et al., 1983); the endosomal shuttling model (Griffiths and Gruenberg, 1991) and the maturation model (Murphy, 1991). The shuttling model is based on preexisting early endosomes with a function as a stationary recycling compartment. From this compartment, carrier vesicles are released to recycle back to the plasma membrane or to interact and fuse with late endosomes (Griffiths and Gruenberg, 1991). The maturation model emphasizes the early endosome as a transient compartment where the transport between early to late endosomes is driven by endosomal maturation. With this model the early endosomes are constantly regenerated through endosomal maturation and are not stationary compartments (Murphy, 1991). Later shuttling models have also shown that the carrier vesicles released by the sorting early endosome goes through maturation trafficking between the early and late endosomes (Scott et al., 2014). The early endosomal maturation process involves changes in the membrane markers as well as the lipid composition of the endosomes (Ebner et al., 2019; Huotari and Helenius, 2011). This pivotal moment in the endocytic pathway is crucial for the sorting of internalized macromolecules and receptors, and it is regulated through an exchange between the two GTPases, Rab5 and Rab7a in the endocytic pathway (Rink et al., 2005). EEs are characterized by the presence of the small GTPase Rab5 on the membrane, responsible for recruiting several effectors that determine the identity and functionality of this compartment (Behnia and Munro, 2005; Huotari and Helenius, 2011). The recruitment of Rab5 to the EE membrane occurs after a transition from cytosolic Rab5-GDP to its active, GTP-bound form, promoted by the Rab5 effector Rabex-5 (Barr and Lambright, 2010). Once on the endosome, Rab5 further stimulates its own recruitment in a positive feedback loop (Lippe et al., 2001). The EE will then change characteristics, becoming increasingly acidic and initiate the formation of intralumenal vesicles, retaining cargo destined for lysosomal degradation (Scott et al., 2014). One crucial step in this progression is the conversion from an EE to a LE through maturation, where the LE is formed by an exchange of GTPases, Rab5 for Rab7a (Rink et al., 2005). The replacement of Rab5 for Rab7a is orchestrated by SAND-1/Mon-1 and Ccz-1 and these two cytosolic factors will bind Rab5-GTP on the membrane of the EE and initiate the recruitment of Rab7a, while promoting the dissociation of Rabex-5 from the endosome, leading to the detachment of Rab5 (Poteryaev et al., 2010). Simultaneously, hVps39p, a subunit of the class C VPS/HOPS complex and a putative guanosine exchange factor (GEF) for Rab7a, will promote the transition from Rab7a-GDP to Rab7a-GTP, in a Rab5-dependent manner (Rink et al., 2005). In parallel, a GTPase activating protein (GAP) will induce the Rab5 hydrolysis of GTP to GDP, further contributing to the displacement of Rab5 from the endosome. Once this step of maturation is concluded, the endosome has become a Rab7a positive late endosome that will mature and initiate lysosomal degradation (Hesketh et al., 2018; Vanlandingham and Ceresa, 2009).

In our experiments we have analyzed enlarged endosomes in MDCK cells expressing Invariant Chain (MDCK-Ii) and endosomes in Hela cells. The endosomal enlargement was achieved by expression of the major histocompatibility complex (MHC) class II-associated invariant chain (Ii) (Nordeng et al., 2002; Stang and Bakke, 1997). The cytosolic tail of Ii has fusogenic properties that induce an enlargement of endosomes (Engering et al., 1998; Nordeng et al., 2002; Romagnoli et al., 1993; Stang and Bakke, 1997), without blocking maturation, or the transition from early to late endosome (Engering et al., 1998; Nordeng et al., 2002; Romagnoli et al., 1993; Stang and Bakke, 1997). This effect of Ii has been previously used to enlarge and study endosomal fusion/fission, coat binding kinetics, morphology, and maturation (Bergeland et al., 2008; Haugen et al., 2017; Landsverk et al., 2011; Skjeldal et al., 2012).

The RabGTPase cycle has become the standard to define endosomal progression and maturation (Rink et al., 2005). However, performing live cell imaging, we were able to provide new data on endosomal maturation and progression through Rab5 to Rab7a exchange. Rab5 detachment from the maturing endosome occurs in a stepwise manner where the initial detachment is independent of Rab7a, whereas the completion of the Rab5 detachment depends on endosomal Rab7a recruitment (Chotard et al., 2010; Li et al., 2009). During the completion phase Rab5 converges to a “hot spot” on the endosomal membrane where the concentrated Rab5 acts as a physical cue for the formation of a new Rab5 positive endosome.

This previously undocumented formation of a native Rab5 positive endosome from a “mother” endosome in transition could explain how the homeostasis of the endocytic pool is preserved through the biogenesis of early endosomes during the Rab5/Rab7a exchange.

## Results

### The Rab5 detachment occurs in two phases

Endosomal progression, from early to late endosomes, is carefully regulated by the transition from a Rab5 to a Rab7a positive endosome (Poteryaev et al., 2010; Rink et al., 2005). To better understand this pivotal transition, we performed a live cell study of this particular conversion on endosomes in MDCK-Ii cells and Hela cells (Nordeng et al., 2002; Stang and Bakke, 1997). In MDCK-Ii cells with enlarged endosomes, mCh-Rab5 remains on the endosomal membrane for a longer time (Landsverk et al., 2011). This prolongs the lifetime of an early endosome without changing the actual transition from Rab5-positive EEs to Rab7a-positive LEs (Landsverk et al., 2011). MDCK-Ii cells were transfected with mCh-Rab5 alone (control) or together with EGFP-Rab7a wild type (WT) or the constitutively active mutant EGFP-Rab7aQ67L (Figure 1). During early endosomal progression we observed numerous homotypic fusion events (Skjeldal et al., 2012) where the endosomes grew larger, preparing for the transition from an early to late endosome. Before the transition from Rab5 positive to Rab7a positive endosomes we observed that the early endosomal homotypic fusion events paused. Once the early endosomal homotypic fusion events ceased, we could initiate the single endosomal analysis of the mCh-Rab5 coat detachment and concomitant acquisition of EGFP-Rab7a (Fig 1A, B, C).

**Figure 1.**
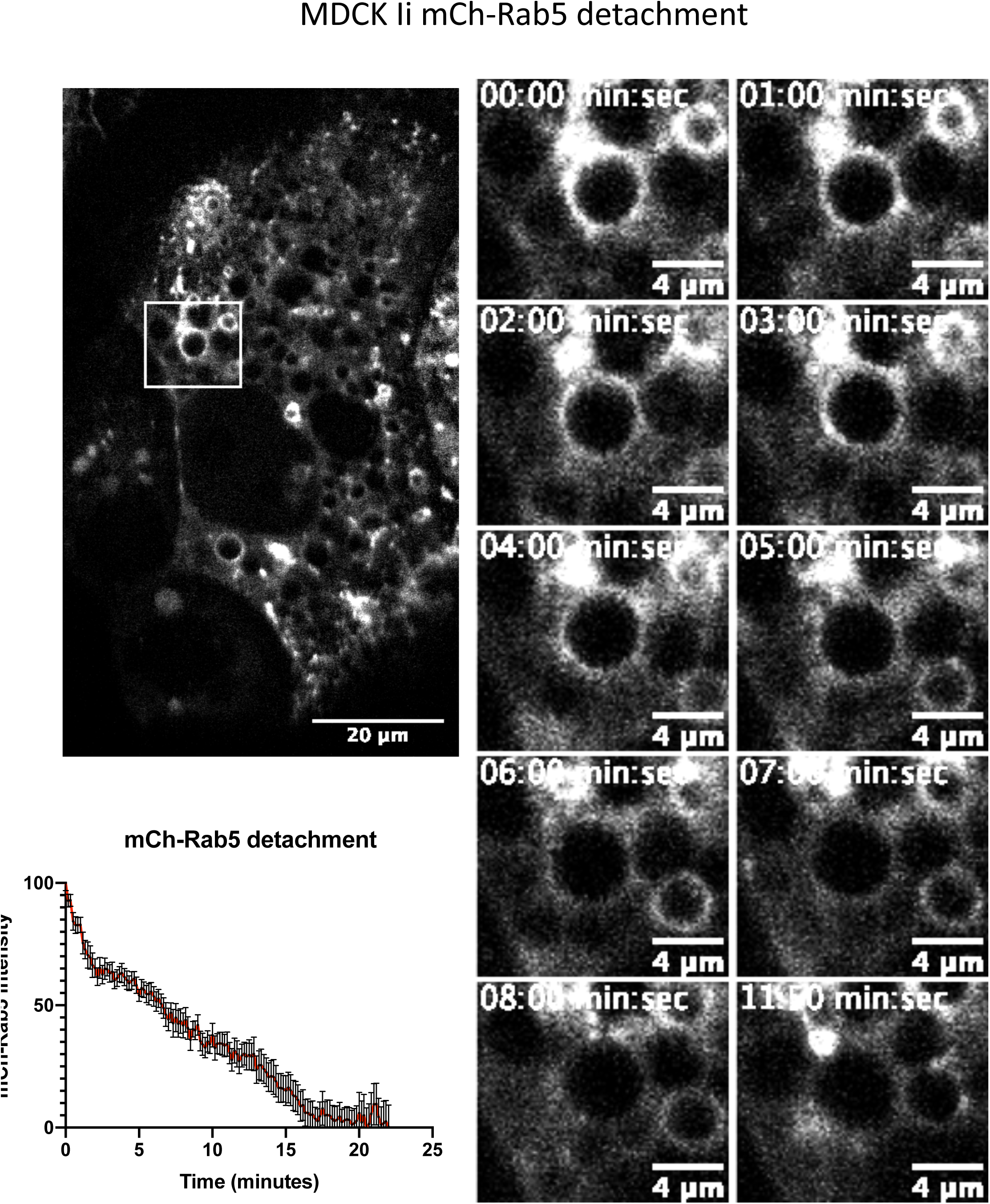
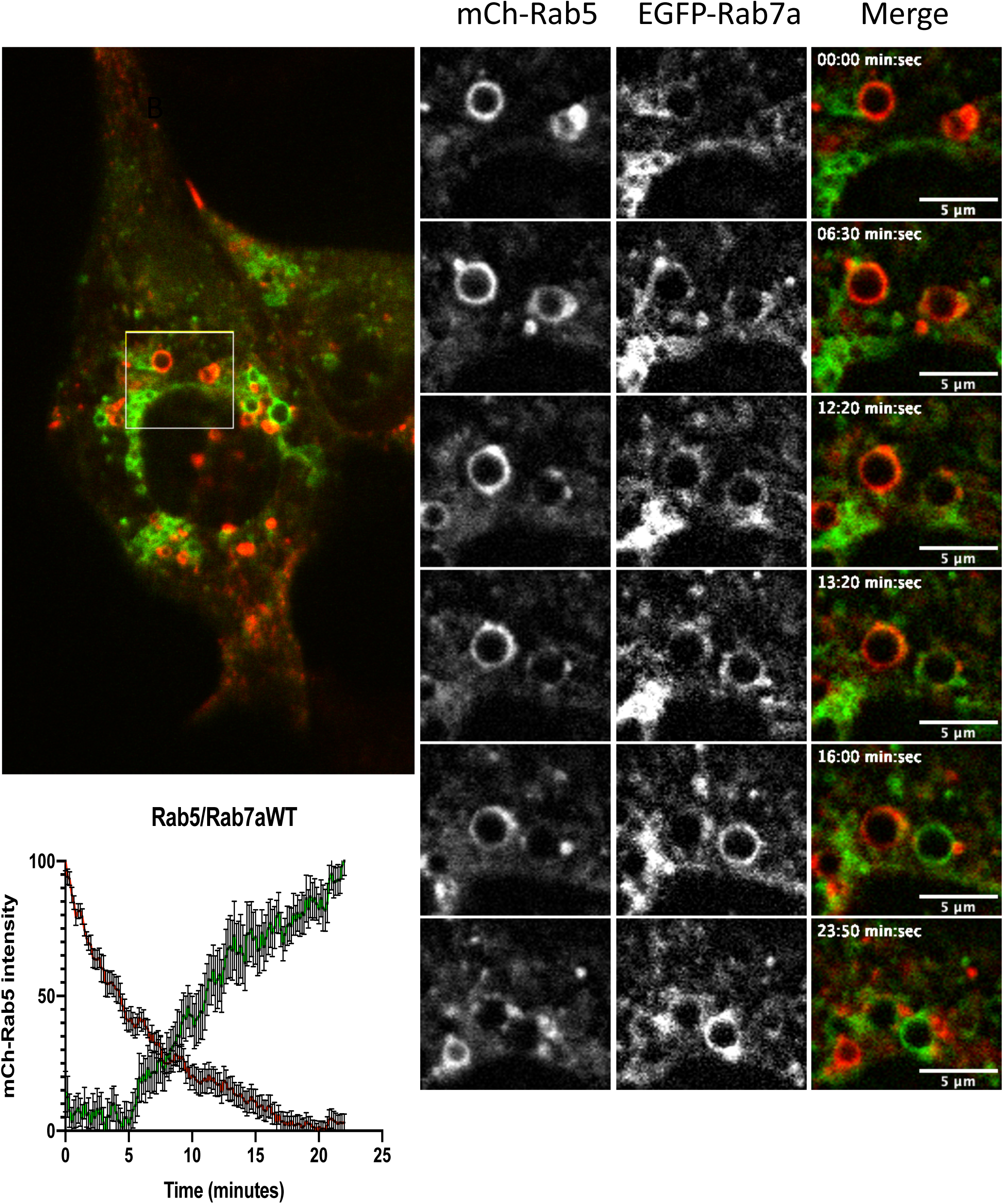
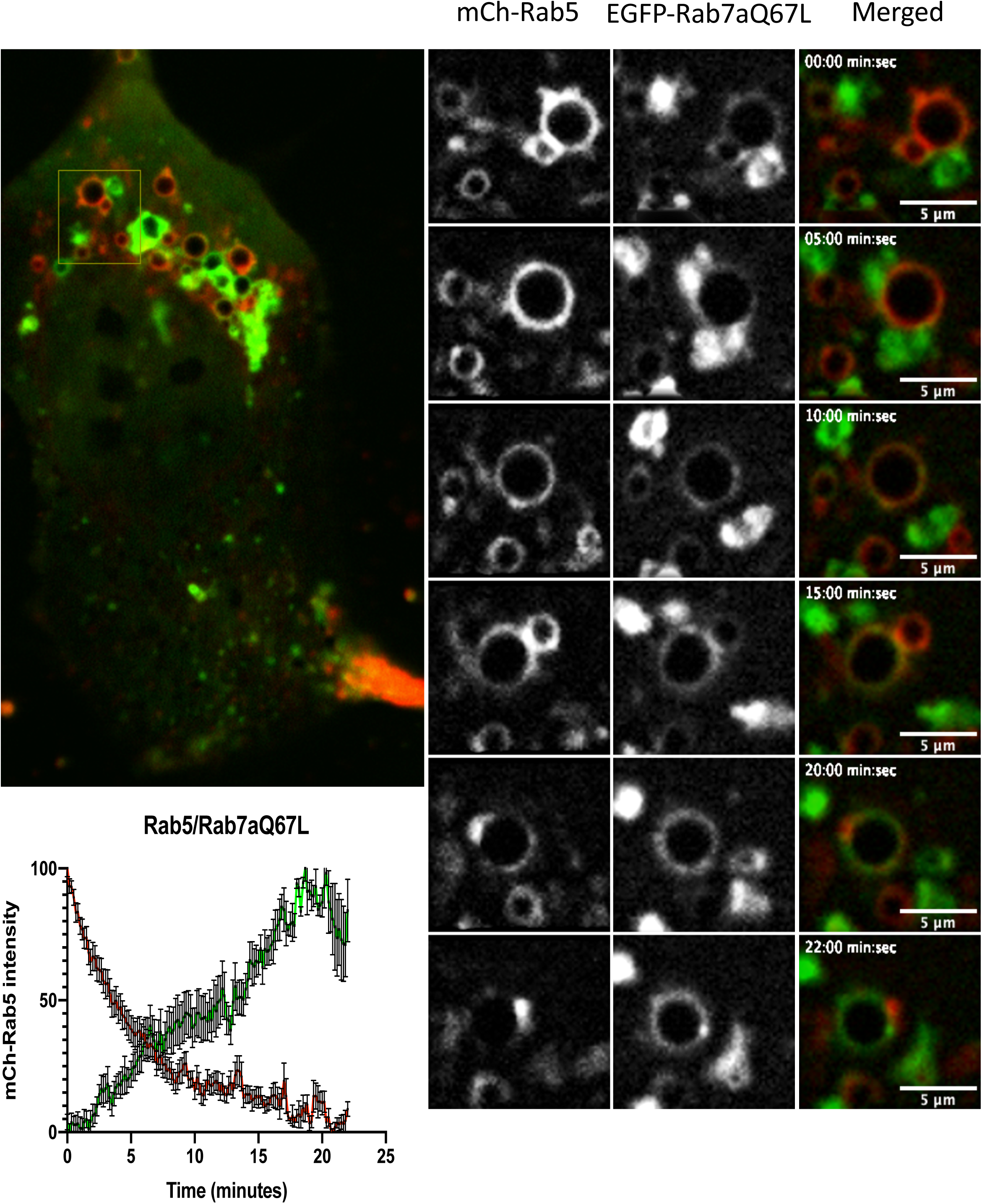
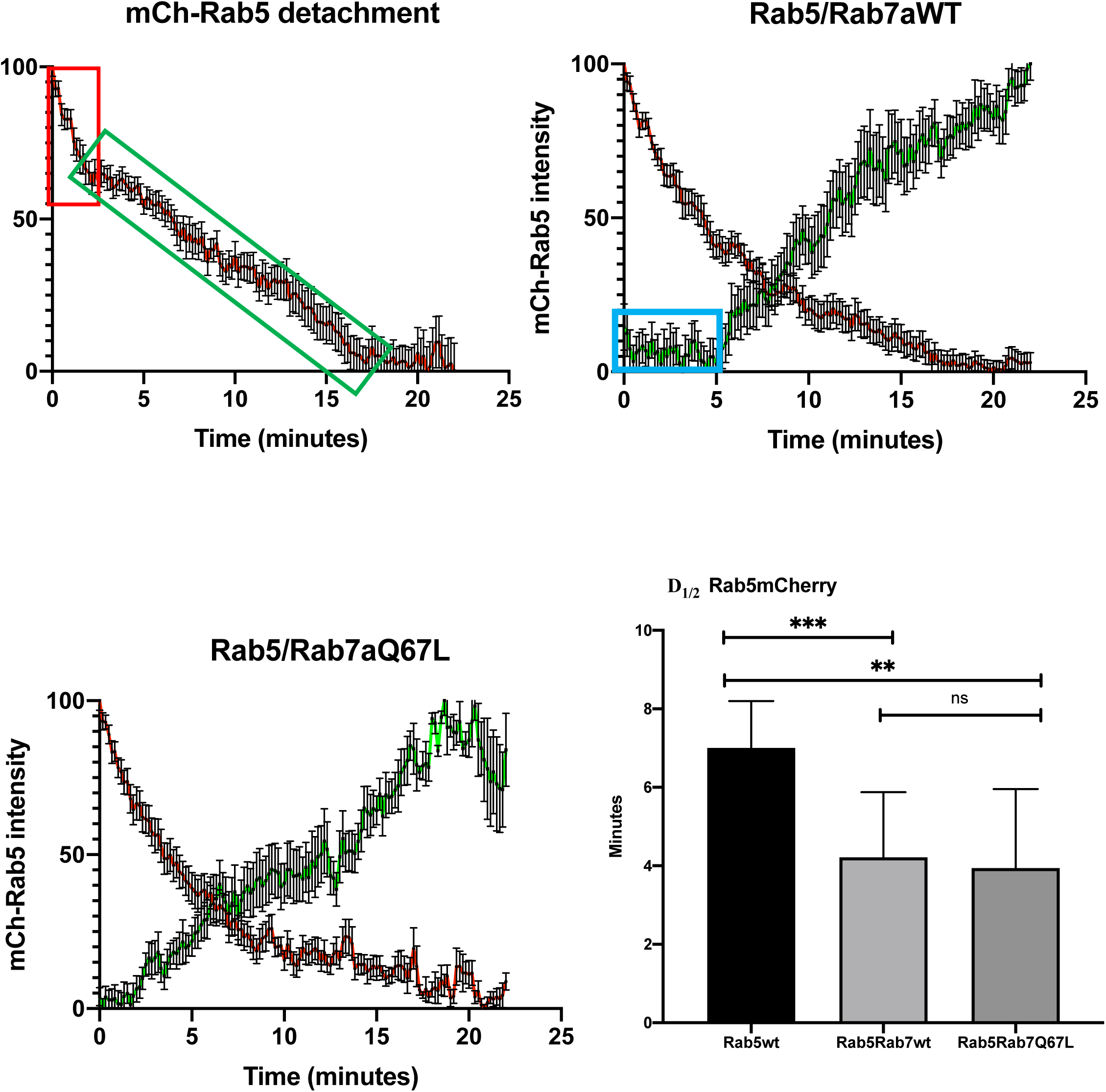
The Rab5 detachment occurs in two phases. A) MDCK cells, stably transfected with invariant chain (Ii) under the control of an inducible metallothionein promoter, were treated with 25µM CdCl_2_ overnight in order to activate the expression of Ii and endosome enlargement. In parallel, the cells were transfected with mCh-Rab5. The graph shows the analysis of mCh-Rab5 detachment on 14 single mCh-Rab5 positive endosomes, averaged over time. Mean = 7.005 min ±, 1.195 min SD, N=14 B) MDCK-Ii cells, stably transfected with invariant chain (Ii) and co-transfected with mCh-Rab5 and EGFP-Rab7a. The graph shows the analysis of mCh-Rab5 detachment on 14 single positive endosomes, averaged over time. Mean = 4.216 min ± 1.659 min SD, N=14. C) MDCK-Ii cells transfected with mCh-Rab5 and EGFP-Rab7aQ67L. The graph shows the analysis of mCh-Rab5 detachment on 14 single positive endosomes, averaged over time., Mean = 3.865 min ± 1.958 min SD, N=14 D) Histogram showing mCh-Rab5 detachment halftime (D_1/2_), 50% drop in intensity from initial value. D_1/2_ was quantified by nonlinear regression (Prism) and averaged over 14 measurements. The analysis shows a significant difference in mCh-Rab5 D_1/2_ from the control experiment and when EGFP-Rab7a or EGFP-Rab7aQ67L is expressed. Significance was tested with a paired t-test between: mCh-Rab5 control to EGFP-Rab7a with p<0.0007 and mCh-Rab5 control to EGFP-Rab7aQ67L with p<0.0012

In control cells (MDCK-Ii transfected with mCh-Rab5), we followed single mCh-Rab5 positive endosomes and plotted the detachment curve (Fig 1A, Movie 1A). Similarly to previously observed Rab5 to Rab7a conversion (Rink et al., 2005) the coat gradually detached from the maturing endosome and became mCh-Rab5 negative (Figure 1A). In order to better understand the mCh-Rab5 coat dynamics during maturation we measured the total time of detachment and calculated the halftime coat detachment (D_1/2_) (see materials and methods). In the control cells (Figure 1A), we measured the D_1/2_ for mCh-Rab5 to be 7.0±1.2 minutes (Figure 1D). To further analyze if a higher expression level of Rab7a could affect the endosomal Rab5 detachment, we double-transfected MDCK-Ii cells with mCh-Rab5 and EGFP-Rab7aWT (Figure 1B, Movie 1B). As previously shown (Rink et al., 2005) we could observe a gradual exchange between Rab5 and Rab7a (Figure 1). Analysis further showed that a higher expression of Rab7a induced a faster D_1/2_ for mCh-Rab5, 4.2±1.7 minutes (Figure 1B). We could additionally measure a similar effect on the mCh-Rab5 D_1/2_ when we expressed the GTP bound form of Rab7a, EGFP-Rab7aQ67L, the mCh-Rab5 D_1/2_ was 3.9±1.9 minutes (Figure 1 C, Movie 1C), indicating that Rab7a recruited to the endosomal membrane might regulate Rab5 detachment during endosomal conversion. Comparing the D_1/2_ between the three experiments we could measure a significant difference in the D_1/2_ for mCh-Rab5 when the cells were transfected with either of the Rab7a constructs (Figure 1D).

Further analysis of the characteristics of the detachment graph in the control experiment revealed a new pattern of endosomal coat dynamics. We could clearly measure two different phases for the mCh-Rab5 detachment; an initial fast detachment the first 2-3 minutes, (red box, Figure 1D) followed by a slower phase, here termed the completion phase, where the curve is less steep (green box Figure 1D). In another set of experiments where the cells were transfected with EGFP-Rab7a WT we could measure a lag period before Rab7a got recruited to the endosome (blue box figure 1D) coinciding with the Rab5 fast and initial detachment. In fact, 50% of Rab5 detached before any Rab7a was detected on the endosomes. However, when the cells were expressing the GTP bound form of Rab7a, EGFP-Rab7aQ67L, the lag period was not detected.

These experiments show us that the mCh-Rab5 detachment dynamics can be divided in two phases, an initial fast phase and a slower, completion phase. In addition, these experiments suggest that the recruitment of Rab7a to the maturing endosome occurs only after the initial detachment of Rab5, and that Rab7a may be important for the slower completion phase, as illustrated by the fact that expression of Rab7a increase the D_1/2_ Rab5 detachment rate.

### Initial Rab5 detachment is Rab7a independent

We find that the endosomal maturation starts with a fast detachment of Rab5 before any Rab7a is detected on the membrane (Figure 1). Our analysis showed that the onset of Rab7a attachment to the endosomes is occurring after Rab5 has started to detach (red and blue box fig 1D) at a time where ∼50% of Rab5 has already left the endosome. To further address the significance of Rab7a recruitment for the dissociation of mCh-Rab5 from the endosome, we co-transfected mCh-Rab5 with the dominant negative mutant of Rab7a (EGFP-Rab7aT22N) or depleted Rab7a with specific siRNA (siRab7a) in HeLa and MDCK-Ii cells (Figure 2). Rab7aT22N is locked in the GDP state and thus localized in the cytosol, acting as a dominant negative mutant of Rab7a (Bucci et al., 2000; Feng et al., 1995).

**Figure 2.**
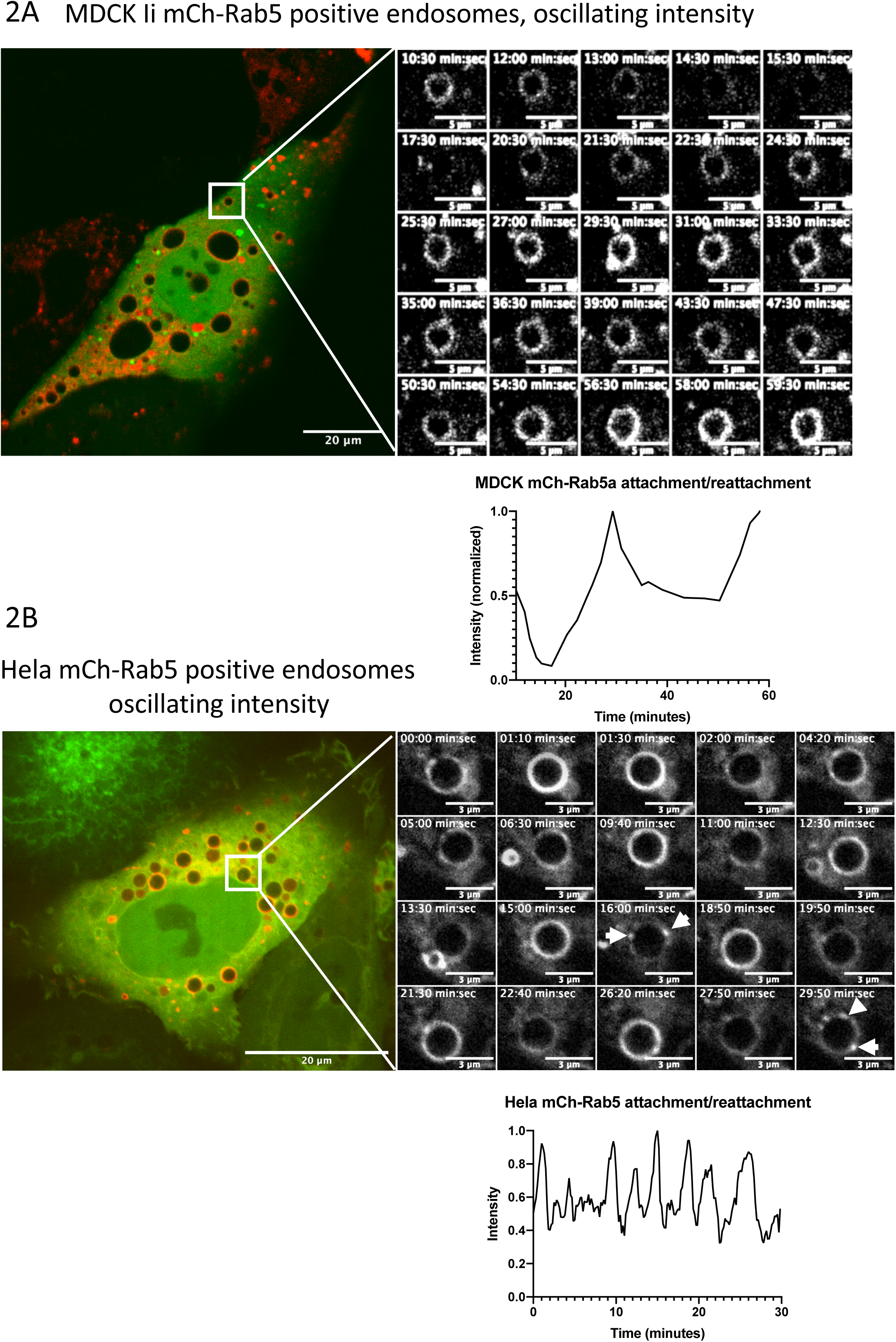
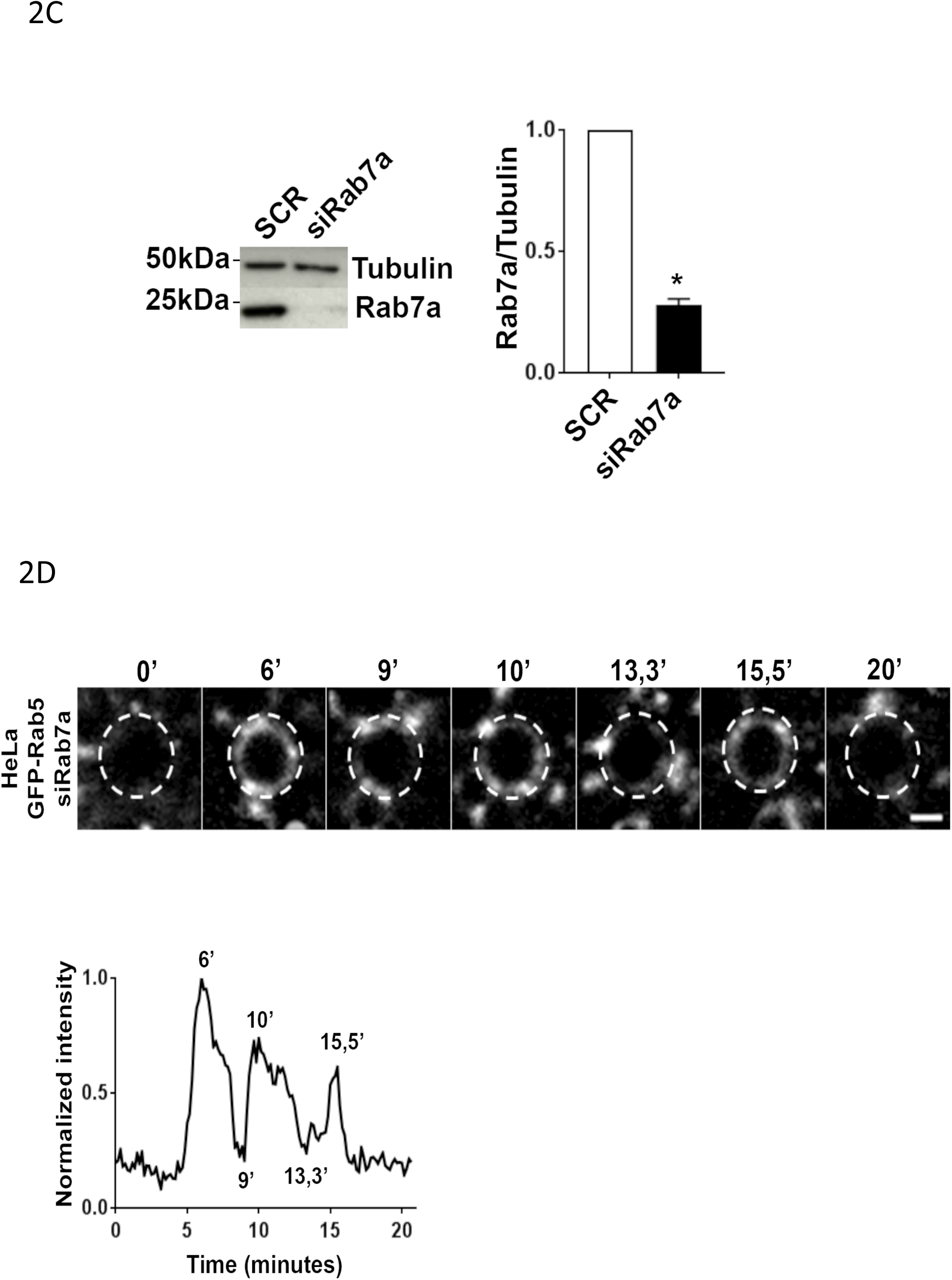
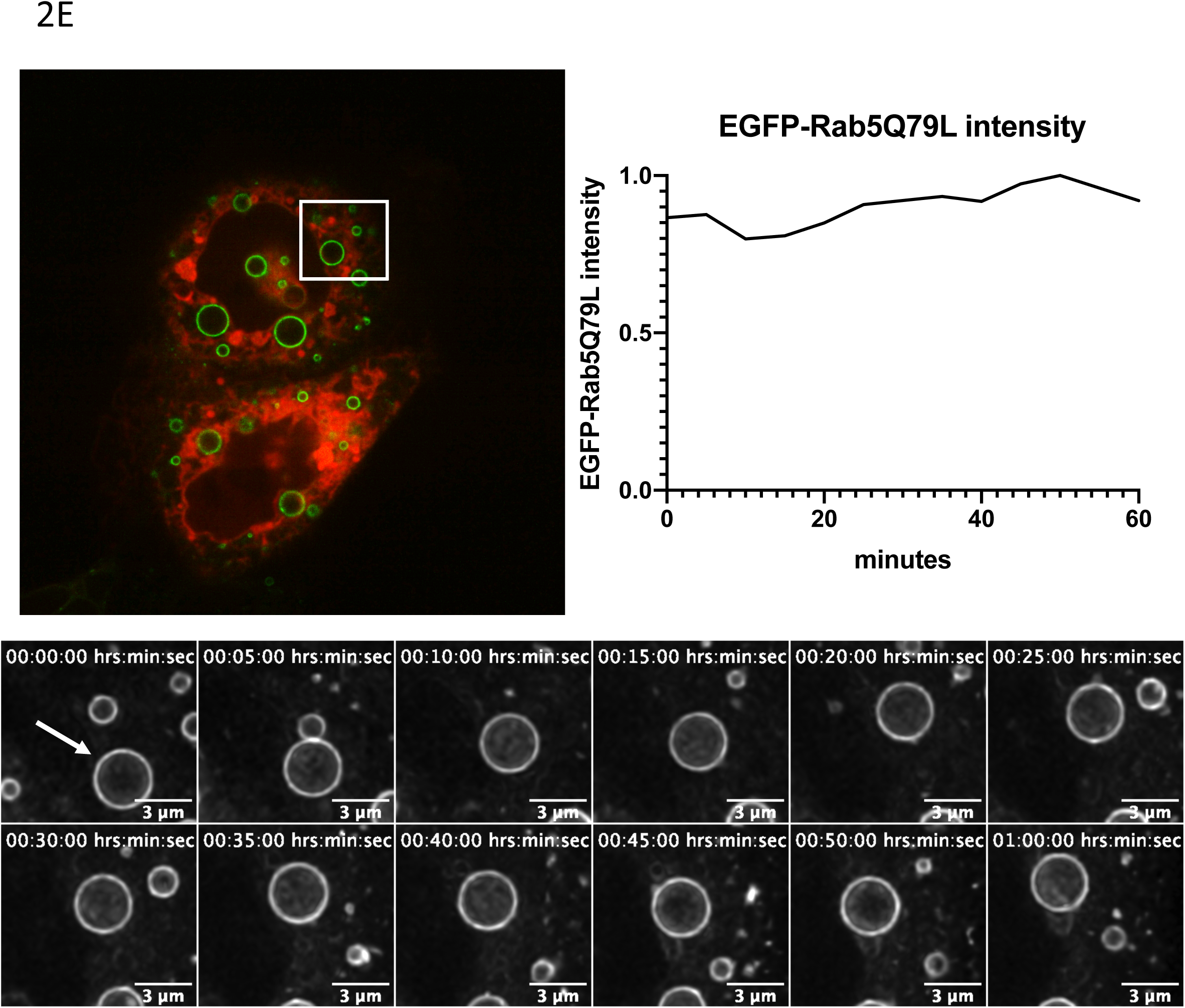
Initial Rab5 detachment is Rab7 independent. A) MDCK expressing Ii induced enlarged endosomes double transfected with mCh-Rab5 and the dominant negative mutant, EGFP-Rab7aT22N. The series of images show detachment and reattachment of mCh-Rab5 induced by the dominant negative mutant. Graphic presentation of the representative mCh-Rab5 detachment. B) Hela cells transfected with mCh-Rab5 and EGFP-Rab7aT22N showing detachment and reattachment with a higher frequency than in similar experiment performed in MDCK. Graphic presentation of the representative mCh-Rab5 detachment. The timestamp on the images are given in minutes. C) Whole cell lysate from SCR and siRab7a in Hela cells were separated by SDS-PAGE and immunoblotted for the indicated proteins. The intensities of the bands were quantified by densiometric analysis and plotted relative to the SCR as mean ± SEM, n=3. D) EGFP-Rab5 positive endosome showing a typical detachment and reattachment induced by the knock down of Rab7a, Scalebar 1μm. The graph represents the normalized intensity of the endosome depicted in the montage. E) EGFP-Rab5Q79L positive endosome in Hela cell transfected with the dominant negative DsRed-Rab7aT22N (Whole cell image). The endosomes in these cells did not show the typical detachment/reattachment pattern of Rab5. Graph showing the fluorescent intensity of EGFP-Rab5Q79L on the representative endosome indicated with a white arrow.

In MDCK-Ii cells co-transfected with mCh-Rab5 and EGFP-Rab7aT22N (Fig. 2A) we observed that the early endosomes remained Rab5 positive for a longer time period and that mCh-Rab5 did not completely detach, on the contrary, mCh-Rab5 detached and reattached, never to fully complete the detachment (Figure 2A). Performing the same experiment in Hela cells we could observe an increase in size of the mCh-Rab5 positive endosomes in EGFP-Rab7aT22N transfected Hela cells (Figure 2 B, Movie 2B). Single endosomal analysis showed a similar and repetitive cyclic pattern, detachment and reattachment of mCh-Rab5 and with a higher frequency (Figure 2 B) The frequency seemed to be quite stable for the endosomes in each cell and the amplitude between the maximum mCh-Rab5 intensity varied between 2-4 minutes (2B graph). Furthermore, depletion of Rab7a using RNAi in Hela cells showed a similar partial mCh-Rab5 detachment and the reattachment (Figure 2C, D). We could observe from the graphs that the dissociation of mCh-Rab5 was fast and seemed to be similar to the fast-initial phase of the mCh-Rab5 detachment (Figure 1D, red box). Careful analysis of the endosomal membrane we could observe mCh-Rab5 positive domains as an intermediate state between the maximum intensity during the intensity oscillations (Figure 2B, white arrows). This shows that the mCh-Rab5 positive endosomes fail to reach the second phase, the completion phase, when no Rab7 is recruited to the endosome in transition.

To search for a role of the Rab5 GDP/GTP cycle in this process we transfected Hela cells with the constitutively active mutant of Rab5 (EGFP-Rab5Q79L), which is unable to hydrolyze GTP and remains membrane-bound (Stenmark et al., 1994), together with DsRed-Rab7aT22N. Hela cells expressing EGFP-Rab5Q79L induced enlarged endosomal structures (Wegner et al., 2010) that had a uniform EGFP-Rab5Q79L positive membrane for more than one hour (Figure 2E). Under these conditions, no cyclic variation was seen for the EGFP-Rab5Q79L positive membrane intensity (Figure 2E). As the Rab7a dominant negative mutant did not induce detachment/reattachment of EGFP-Rab5Q79L, this indicate that the initial fast detachment phase of Rab5 is regulated by the rapid GDP/GTP cycle.

These above results demonstrate that Rab5 has the ability to transiently leave the maturing endosome independently of membrane associated Rab7a, but Rab7a has to be recruited to the endosome membrane for Rab5 to complete its detachment. Overall, these results show that mCh-Rab5 detachment occurs in two phases: the initial phase, which is Rab7a independent; and the completion phase, which is Rab7a dependent.

### Rab5 domains converge and leaves the maturing endosome

During the analysis of single maturation events on Ii-enlarged endosomes in MDCK-Ii cells (Figure 1) transfected with mCh-Rab5, we consistently observed an increase in the formation of endosomal Rab5 positive domains (Figure 1 and 3 A, Arrows). In a typical experiment presented in figure 3A we could follow mCh-Rab5 positive enlarged endosomes changing from having a uniform mCh-Rab5 membrane to acquire several mCh-Rab5 domains. During the Rab transition these domains converge towards one (Figure 3A, Series 1) or two (Figure 3A, Series 2) distinct mCh-Rab5 domains to complete the second phase of maturation, completion phase. Converging mCh-Rab5 domains appeared in all of the measured endosomes going through the steps of Rab5 detachment. Similar experiments were performed with super-resolution imaging (Zeiss AiryScan) on Hela cells transfected with EGFP-Rab5 and mApple-Rab7a, where the converging domains were confirmed on endosomes not enlarged with Ii (Figure 3B, arrows, Movie 3B). Similarly, to the previously measured enlarged endosomes, the formation of EGFP-Rab5 domains on smaller endosomes started after the initial rapid Rab5 decrease and occurred throughout the entire completion phase. Endosomal microdomain formation and convergence to one or two major domains occurred in both Ii enlarged endosomes and in endosomes in Hela cells.

**Figure 3.**
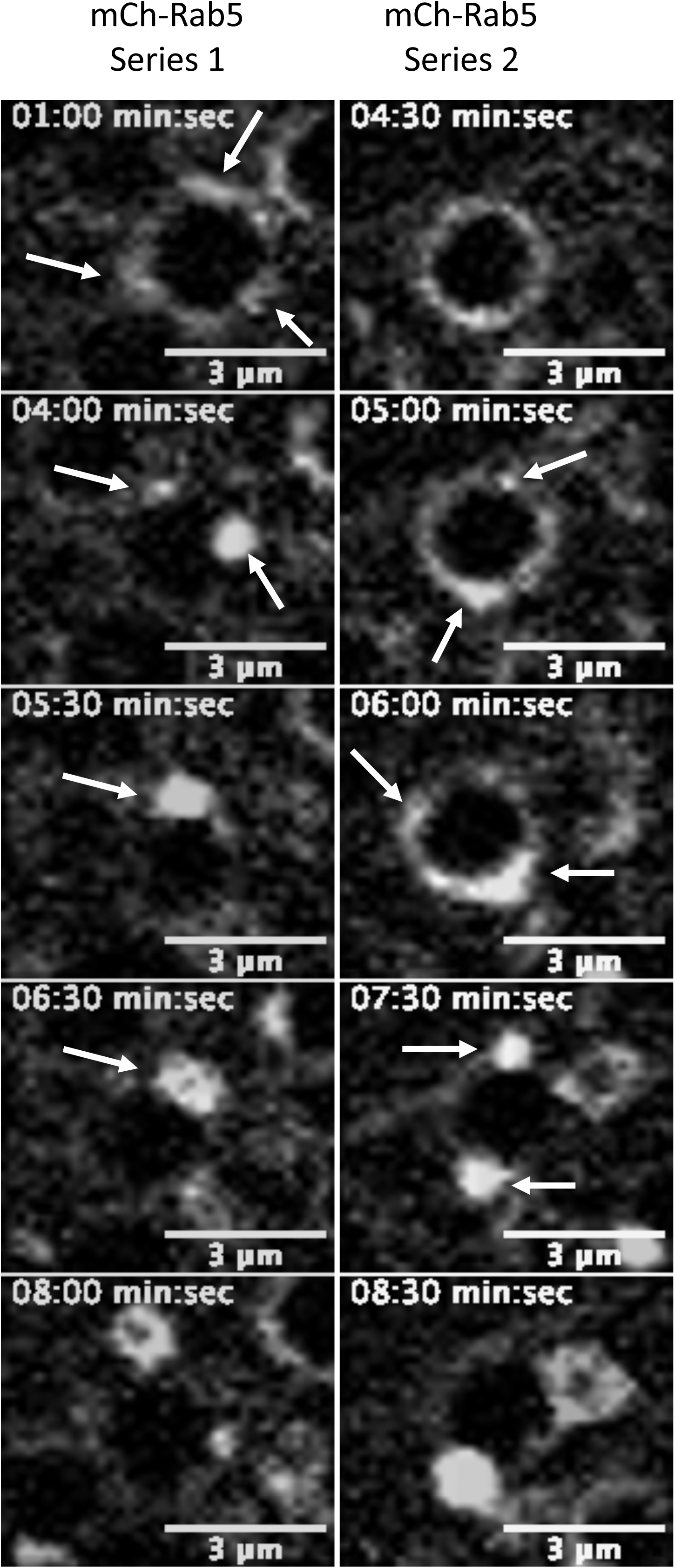
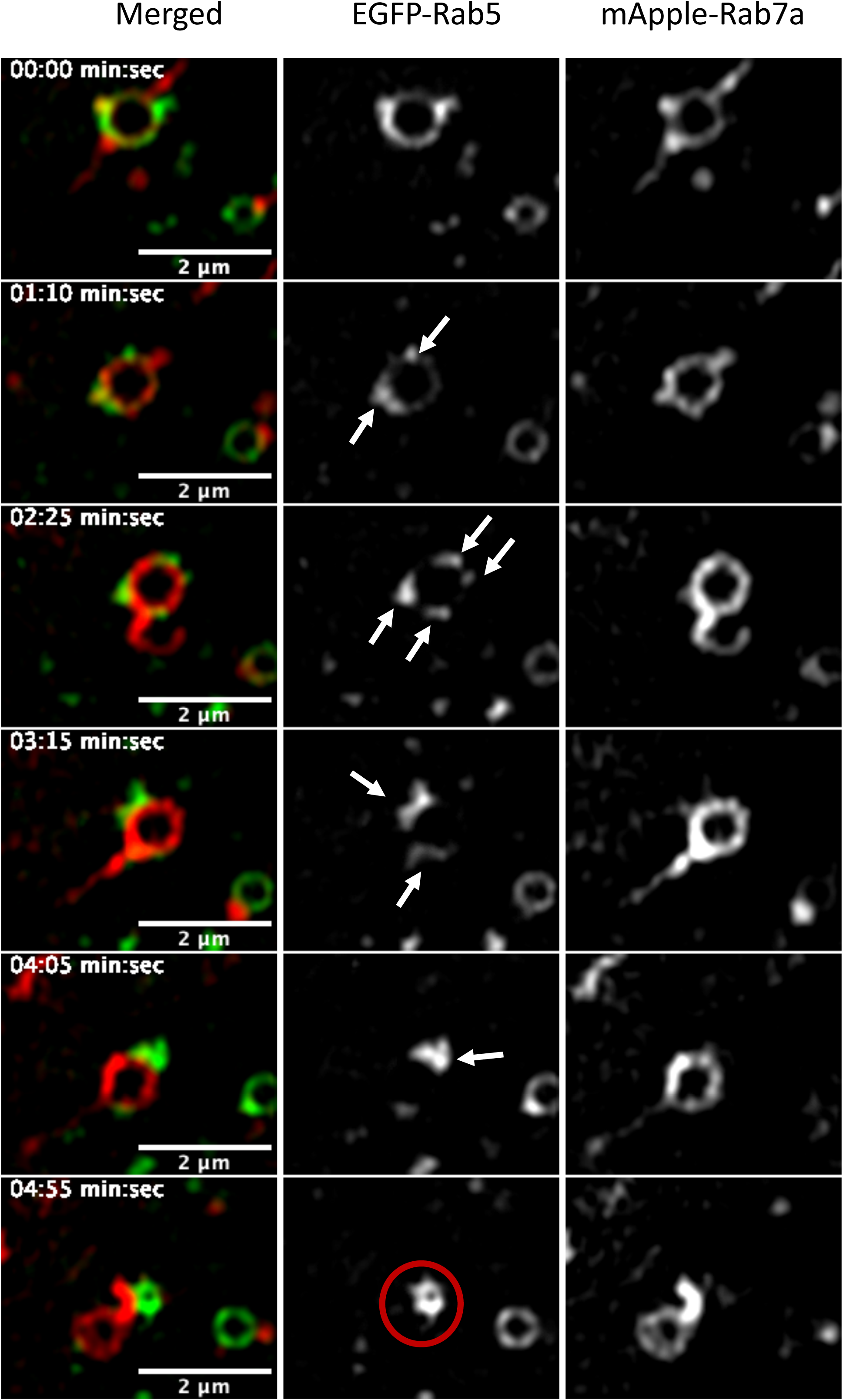
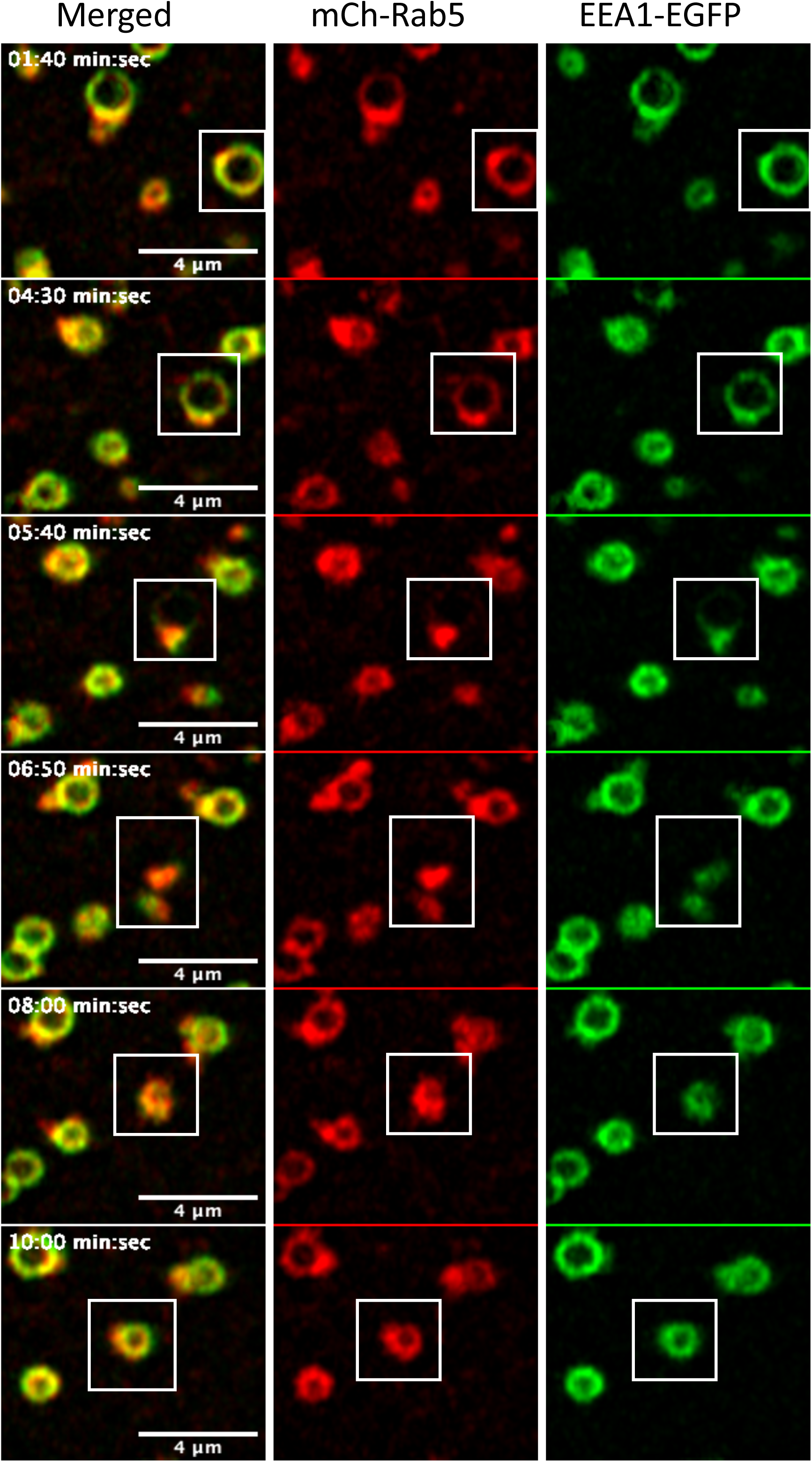
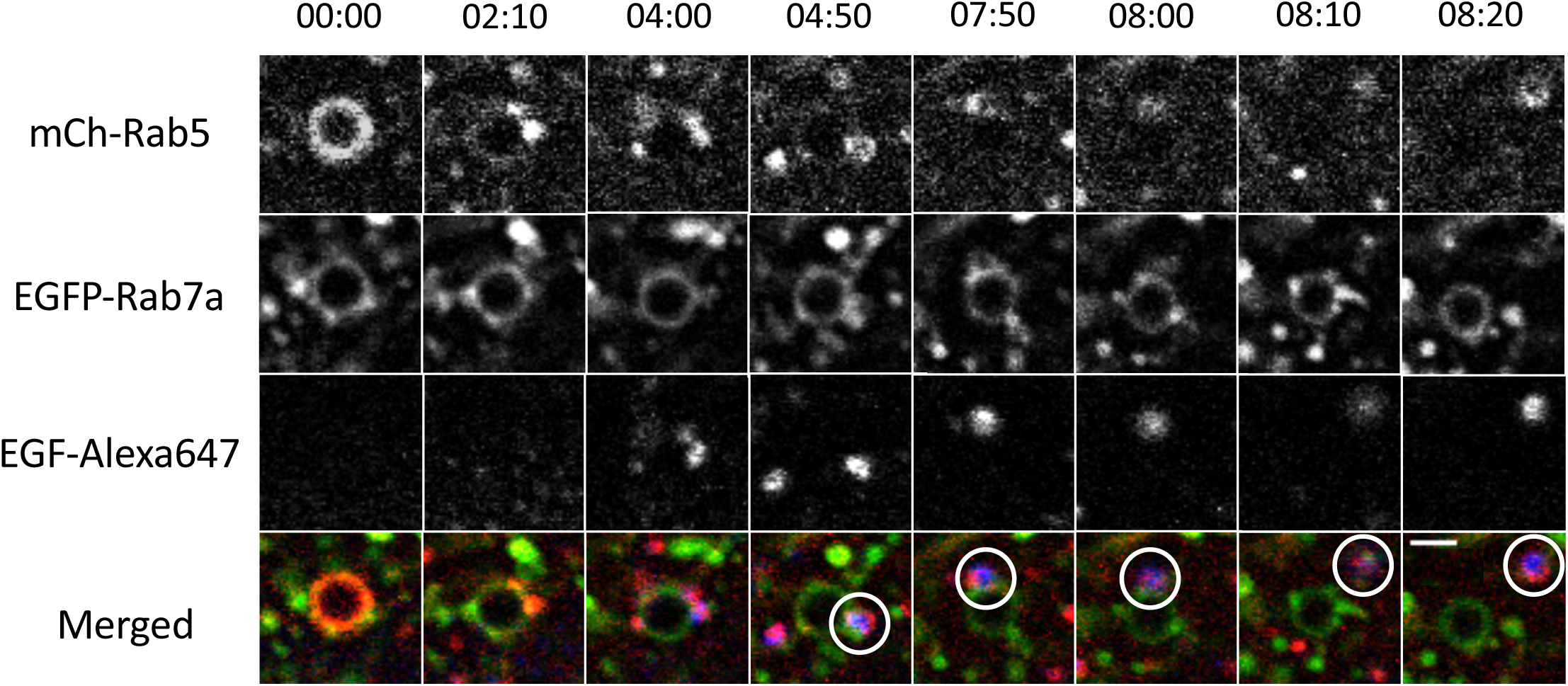
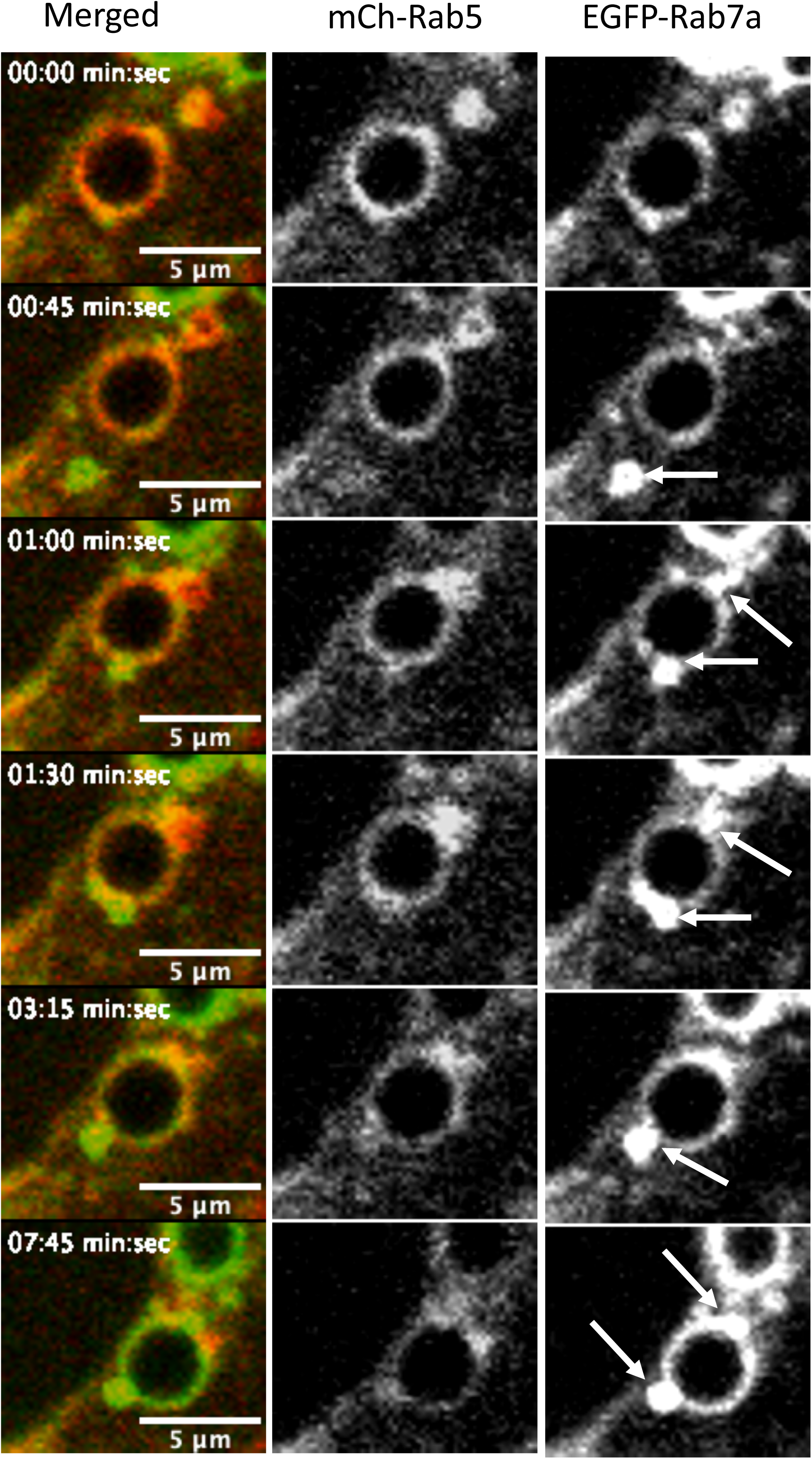
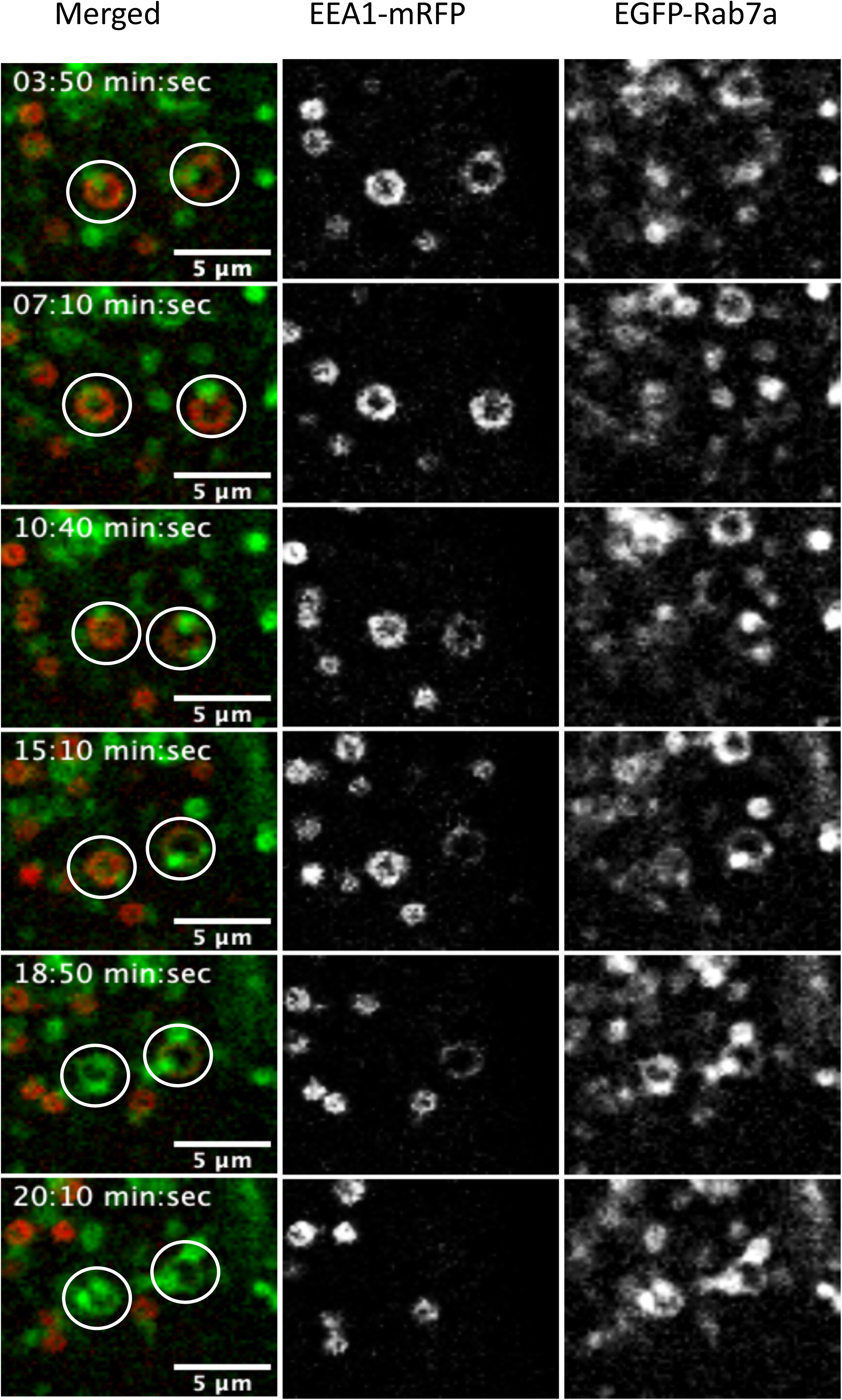
Second phase of endosomal Rab5 to Rab7 transition, Rab5 convergence. A) Two mCh-Rab5 positive endosomes in MDCK-Ii cells (series 1 and 2) going through the convergence phase. Arrows indicate the formation of mCh-Rab5 positive domains that converge to one or two main domains to accomplish the mCh-Rab5 detachment. B) Hela cells expressing EGFP-Rab5 and mApple-Rab7a (::1μm in diameter) positive endosomes going through the convergence phase of Rab5 to Rab7a transition. Arrows indicate the formation of EGFP-Rab5 positive domains converging to one major domain to accomplish the Rab5 detachment through the formation of a (daughter) EGFP-Rab5 positive endosome (red circle). C) Hela cells expressing mCh-Rab5 and EEA1-EGFP wt positive endosomes progressing through convergence phase to give rise to a newly formed mCh-Rab5 and EEA1-EGFP wt positive endosome. D) Hela cells expressing mCh-Rab5 and EGFP-Rab7a positive endosomes during uptake of EGF-Alexa 647. The newly formed mCh-Rab5 and EGFP-Rab7a positive endosome from the maturing endosome acquire EGF-Alexa 647 showing a fully functional newly formed endosome (white circles). Time is represented as min:sec and scalebar is 2µm. E) MDCK-Ii cells transfected with mCh-Rab5 and EGFP-Rab7a showing the gradual contribution of incoming EGFP-Rab7a throughout Rab5 to Rab7a transition. During mCh-Rab5 detachment, EGFP-Rab7a positive endosomes are recruited to the maturing endosome to provide the necessary EGFP-Rab7a. F) MDCK-Ii cells transfected with EEA1-mRFP wt and EGFP-Rab7a showing the same EGFP-Rab7a recruitment to the maturing endosome.

By following the fate of the converging endosomal Rab5 domains, we observed that these domains invariably gave rise to a newly formed mCh-Rab5 or EGFP-Rab5 positive endosome (Figure 3 B, red circle). The Rab5 positive and converged domain leaving the maturing endosome seemed to work as a prerequisite for the formation of a new early Rab5 positive endosome. We then asked if this mechanism of maturation was unique for Rab5. In order to better understand this, we co-transfected cells with a well-known tethering protein EEA1-GFP wt (Bergeland et al., 2008; Murray et al., 2016; Simonsen et al., 1998) together with mCh-Rab5 and analyzed the coat detachment of the respective membrane associated proteins. The detachment of the two membrane associated proteins occurred simultaneously and the convergence towards a local endosomal hotspot was observed with both EEA1-GFP wt and mCh-Rab5. This indicates that both EEA1 and Rab5 follows a similar endosomal detachment pattern (Figure 3C, white box, Movie 3C). Further analysis of the Rab5-EEA1 positive vesicles showed that after their release from the newly matured endosome, they instantly engaged in early homotypic fusion (Figure 3C). EEA1 and Rab5 are kept on the membrane as a fully functional fusion machinery to ensure that the newly released endosomes are instantly fully functional to engage in homotypic fusion. Early endosomes work as sorting stations for protein destined for degradation or recycling (Huotari and Helenius, 2011). In order to better understand the function of the newly formed mCh-Rab5 endosome we added EGF-Alexa647 to the cells. This experiment showed us that the newly formed mCh-Rab5 positive endosome could recruit EGF-Alexa647 containing vesicles showing that the newly formed early endosome is primed for another round of sorting of EGF/EGFR (Figure 3D).

During the formation of mCh-Rab5 positive domains we could observe an increase of EGFP-Rab7a on the maturing endosome in domain like structures (Figure 1). Throughout the Rab5 to Rab7a transition we could observe an increase of EGFP-Rab7a positive vesicles clustering around the endosomes as Rab5 was detaching. These incoming and interacting endosomes seemed to deliver EGFP-Rab7a to the maturing endosome (Figure 3E, Movie 3E). This experiment was repeated with MDCK-Ii cells co-transfected with EEA1-mRFP wt and EGFP-Rab7a showing the same result (Figure 3F, Movie 3F) the endosomes in transition seem to acquire the Rab7a through a fusion like or “kiss and run” process. These incoming EGFP-Rab7a vesicles seem to carry the signal for the maturing endosome to proceed from the initial phase to the convergence part of maturation since in the Rab7aT22N transfected cells the endosomes never reached the completion phase.

### Rab5 converged domains consist of the membrane bound immobile fraction

We have shown that Rab5 detachment occurs in two phases: an initial rapid and a slower convergence phase after Rab7a is recruited to the endosome. Rab GTPases alternates between a membrane bound and a cytosolic state and we can measure the immobile and the mobile fraction present on the endosomes (Sprague and McNally, 2005). We have previously described that endosomal associated proteins can specifically alter the ratio between the mobile and the long lived immobile fraction both as a function of signaling and between interphase and mitosis (Bergeland et al., 2008; Haugen et al., 2017). As this is believed to reflect a functional state, we wanted to investigate whether these two fractions of Rab5 on the endosomal membrane varied through to the two detachment phases of Rab5.

To analyze the binding kinetics of Rab5 before and during conversion, we performed single endosome FRAP experiments on endosomes prior to transition and on converged domains. In these experiments, we used the MDCK-Ii cells expressing enlarged endosomes transfected with EGFP-Rab5. Analysis of bleaching experiments on single endosomes positive for EGFP-Rab5 showed us that EGFP-Rab5 has a mobile fraction of 22% and an immobile fraction of 78% (Figure 4), similar to previously observed values (Haugen et al., 2017). In order to compare the initial Rab5 detachment with the slower convergence maturation we had to specifically bleach the converging domains before the transfer to the newly formed Rab5 positive endosome. Enlarged endosomes together with a fast bleaching module (Andor Dragonfly mosaic) made this technically possible (Figure 4A). Bleaching experiments and analysis of the converging domains revealed an altered binding dynamic of EGFP-Rab5 in the converged domains compared to the binding dynamics prior to domain formation (Figure 4A). We could measure a significant change in the immobile fraction of EGFP-Rab5 in the converging domains, which increased by a factor of 4 (Figure 4B). This result shows a gradual increase of the immobile fraction during the convergence phase prior to a complete detachment. We could further show that this immobile fraction consisted of the GTP bound form of Rab5 by transfecting the cells with GFP-Rab5Q79L and performing the same FRAP experiment (Figure 4B). Comparing the membrane fractions of the converging domain and the GFP-Rab5Q79L we could show that they are exactly the same, strongly indicating that the converging domains and the Rab5 leaving the endosome to form a new endosome consists of the immobile fraction. These experiments proved that the immobile fraction is 80% in the converging domains and that this is the GTP bound form of Rab5 (Figure 4).

**Figure 4.**
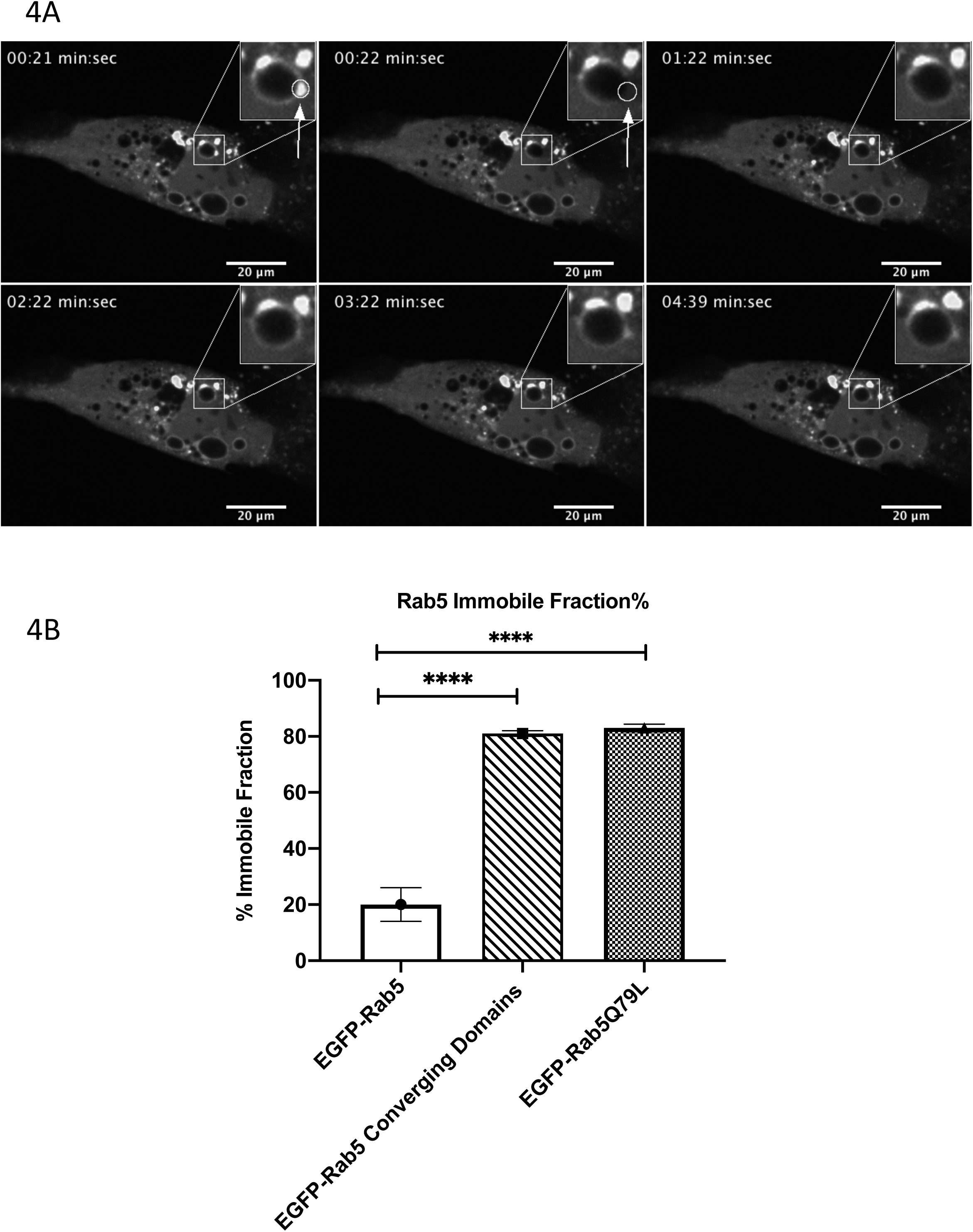
Rab5 converged domains consist of the immobile fraction. A) MDCK-Ii cells transfected with EGFP-Rab5 showing enlarged endosomes positive for EGFP-Rab5 positive domains. One of the EGFP-Rab5 positive domains was bleached and the immobile fraction was calculated. B) Graph showing the immobile fraction of (control) EGFP-Rab5 = 20% ± 6% SD, n=14, immobile fraction EGFP-Rab5 converging domain = 81% ± 1% SD, n=16, immobile fraction EGFP-Rab5Q79L = 83% ± 1.4% SD, n=12.

These results show that Rab5 detaches in two phases; the initial phase may be regulated by the GDP-GTP cycle and pertains to the mobile fraction of Rab5, while the slower phase is a concentration of Rab5 immobile fraction to the converging domains prior to a transfer from the maturing endosome to prime the formation a new early endosome.

## Discussion

Endosomal trafficking is carefully regulated through organelle specific Rab GTPases (Bhuin and Roy, 2014; Zerial and McBride, 2001). These GTPases identify and regulate the progression of endocytosed macromolecules and plasma membrane receptors (Langemeyer et al., 2018). Progression and direction from the early to the late endosome have been shown to be regulated by a Rab5 to Rab7a switch (Cabrera et al., 2014; Huotari and Helenius, 2011; Nordmann et al., 2010; Poteryaev et al., 2010; Rink et al., 2005). This switch/conversion have previously been described entirely as an exchange between the cytosolic pool and the membrane bound fraction of Rab5 and Rab7a, specifically regulated through the Rab GTP/GDP cycle (Pfeffer, 2001; Pfeffer, 2017; Stenmark and Olkkonen, 2001).

In this paper we have shown that enlarged early endosomes in MDCK-Ii cells (Landsverk et al., 2011; Skjeldal et al., 2012; Stang and Bakke, 1997) and early endosomes in Hela cells, mature into late endosomes by a stepwise Rab5 detachment, controlled but not initiated by Rab7a. We show that the detachment of Rab5 follows a two-step mechanism where the initial detachment is fast and diffusion-like, as expected for the GTP/GDP cycle, and the second part is slower and regulated by converging Rab5 domains (Figure 1). When Rab5 were co-transfected with Rab7a, either wt or Q67L mutant, we could measure a faster D_1/2_ of Rab5, indicating a Rab7a regulatory role of Rab5 detachment, specifically in the converging phase of Rab5 detachment. Furthermore, Rab7a was not recruited to the maturing endosome before the initial phase was completed, as shown in the Rab7a lag phase (Figure 1D, blue box). This indicates that the Rab5 detachment has a Rab7a independent phase and a Rab7a dependent phase, which was confirmed after co-transfection with Rab7aT22N (dominant negative mutant) or knocking down Rab7a with RNAi. Under these conditions the full detachment of Rab5 did not occur and the fluorescent endosomes initiated an association/disassociation like characteristic of Rab5 coat detachment (Figure 2). Apparently, Rab5 detachment could not be completed because Rab7a positive vesicles were not recruited to the endosome to initiate the converging phase of Rab5 detachment. When we expressed the constitutively active Rab5 together with the dominant negative Rab7a mutant this specific “blinking” characteristics did not occur, indicating that the initial detachment of Rab5 is regulated by the activity of its GAP, independently of any possible feedback-loops later established by Rab7a.

The show here that the feedback loop from the endosomal system consists of Rab7a positive endosomes being recruited to the Rab5 maturing endosome after the initial and fast Rab5 detachment. In the experiments with the dominant negative Rab7a mutant or Rab7a knockdown, where there were no “incoming” Rab7a positive endosomes, the Rab5 positive endosomes were not able to proceed from the GTP/GDP regulated phase to the converging phase, and consequently the incomplete Rab5 association/disassociation pattern could be observed. In between the mCh-Rab5 maximum intensity peaks we could observe the formation of specific domains that were abruptly terminated when the uniform mCh-Rab5 returned. It seems like the endosomes tries to reach the converging phase but lack the signal, which is Rab7 positive vesicles interacting with the maturing endosome. The “incoming” Rab7 positive endosomes provides Rab7 for the maturing endosome and concomitantly transmit the signal to enter phase number two, the converging phase.

Following the converging phase of Rab5 domains on the endosome we could show that the domains would aggregate in one or two domains. This Rab5 positive domain remained bound to the membrane for a longer time period before detaching from the maturing endosome and giving rise to a newly formed Rab5 positive endosome. Recycling of Rab5 through the transfer and formation of Rab5 to a newly formed endosome has previously been discussed to be an alternate mechanism additional to the Rab-GDI mediated exchange (Rink et al., 2005). Here we provide evidence for a direct Rab5 transfer from the maturing endosome to form a novel Rab5 positive endosome.

We could furthermore show that the Rab5 effector protein EEA1 did also follow Rab5 during convergence and transferred together with Rab5 onto a newly formed endosome. The released endosome engaged in early endosomal fusion immediately post formation and could recruit endocytosed EGF, showing that it is a fully functional early endosome ready for another round of maturation.

To better understand the binding dynamics of Rab5 during the convergence phase we performed FRAP experiments on Rab5 positive endosomes before convergence and correspondingly bleached the converging domains. We could show that a Rab5 positive endosome, before the conversion, had 20% immobile fraction and 80% mobile fraction as previously published (Sann et al., 2012). However, in converged domains we measured a total change in the fractions, where we found 80% to be immobile and 20% to be mobile. In the control experiment where we bleached the dominant active mutant of Rab5 (EGFP-Rab5Q79L) we found the same distribution between immobile fraction and mobile fraction as in the converged Rab5 positive domain strongly suggesting that the fraction of Rab5 that detaches from the maturing endosome and gives rise to a new endosome is transferred as a membrane bound GTP form.

Both EEA1 and Rab5 are fusogenic membrane associated proteins that will have to go into a quiescent state during the early endosome to late endosome transition. This may be explained with our FRAP experiment where we measure a redistribution of the immobile and mobile fractions of Rab5. The first initial detachment proved to be the mobile fraction that detaches through the regular GDI mediated GTP/GDP cycle. Inactivating Rab5 fusogenic properties through a redistribution of the mobile fraction to immobile fraction may also change the activity of EEA1 inhibiting the entropic collapse which is necessary for tethering prior to fusion (Murray et al., 2016). We hypothesize that the Rab5 fractions on the newly formed endosome will be redistributed back to mobile fraction 80% and immobile fraction 20% in order to reactivate the fusion machinery. A redistribution of the immobile and mobile fraction of Hrs and Eps15 has previously been published to be important for EGFR degradation (Haugen et al., 2017). Similar redistribution of Rab5 was shown in this study which may indicate that a local redistribution of the mobile and immobile fraction on endosomes is a general mechanism to regulate endosomal maturation and progression.

In this paper we describe a new mechanism on the dynamics of Rab5 and Rab7a during endosomal maturation. The Rab5 to Rab7a switch is definitely not an exchange where Rab5 is directly replaced by Rab7a, the Rab5/Rab7a exchange is regulated in distinct steps. The initial diffusive phase of Rab5 detachment may act as a switch in endosomal transition activating the SAND-1/Mon-1/HOPS cascade (Poteryaev et al., 2010). This is to actively recruit the Rab7a positive endosome necessary for completing convergence phase of Rab5 detachment. However, the switch does not seem to control the entire transition but act as a signal to activate the second phase of transition. We find that Rab7a recruitment to the endosome in transition most likely occur through vesicular interactions providing Rab7a to the maturing endosome. We were not able to observe whether the incoming Rab7a positive endosomes fused with the endosome in transition or was transferred from one vesicle to the other by “kiss and run” (Duclos et al., 2003).

Both the shuttling model and the maturation model have contradictory characteristics, especially regarding endosomal homeostasis (Helenius et al., 1983). The shuttle model predicts a set of “pre-existing” endosomal compartments, shuttling carrier vesicles between the early and late compartments (Griffiths and Gruenberg, 1991). To put the “pre-existing endosomes in to context we have to compare the condition before and after mitosis. During cell division the mitotic machinery ensures that the number of endosomal vesicles in the two daughter cells have approximately half the copy number each (Bergeland et al., 2001) and among the sustained vesicles after mitosis we would find the “pre-existing” endosomes. In this study we found that, after mitosis the number of early endosomal vesicles increase the first three hours of the G1 phase and subsequently we observed an increase in the number late endocytic compartments, indicating that the early endosome pool mature to late endosomes (Bergeland et al., 2001). This indicate that all the early endosomes in the daughter cells are “pre-existing” (Kamentseva et al., 2020) and will mature to produce new late endosomes. Consequently, Rab5 convergence induce formation of early endosomes to maintain the homeostasis between early and late endosomes.

We here show that early endosomes through Rab5 to Rab7a switch, continuously give rise to new functional endosomes, a crucial step to maintain the homeostasis of early endosomes. This maturation regulated by transition from early to late endosomes provides the pool of early endosomes that exert the role as a sorting organelle. Furthermore, the maturation through conversion gives a direction for the endosomal progression. With these novel findings we here present a crucial step in understanding how endosomal trafficking is regulated to maintain endosomal homeostasis and how membrane associated proteins could be recycled during endosomal maturation (See graphical model figure 5).

**Figure 5.**
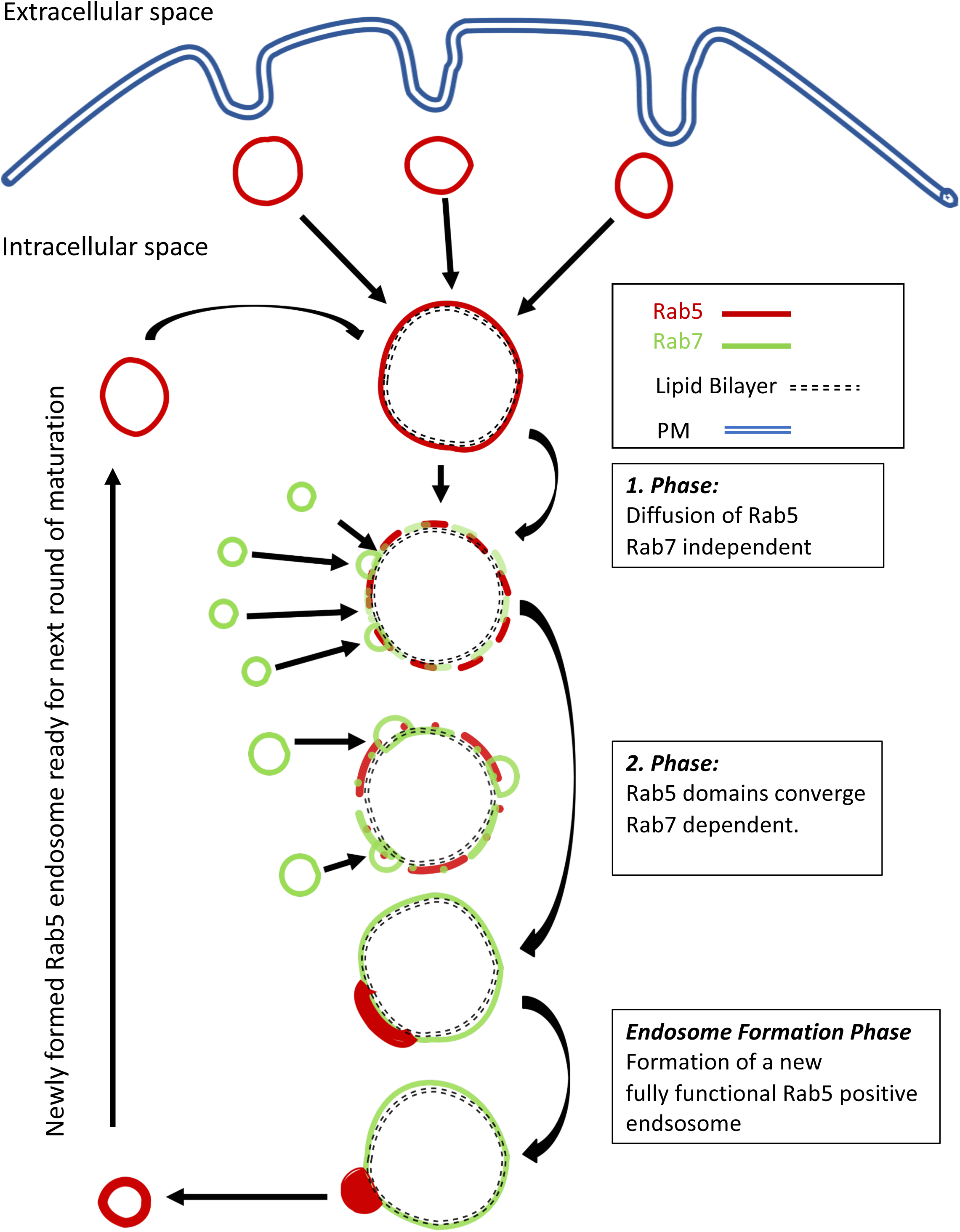
Graphical Model: Endosome maturation to endosome formation model. A model describing the stepwise maturation pattern that give rise to novel Rab5 positive vesicles. Newly internalized Rab5 positive endosomes fuse with themselves and with preexisting Rab5 positive endosomes. Maturation from Rab5 to Rab7a positive endosomes takes place in two phases; 1. Fast initial and diffusion like Rab5 detachment, Rab7 independent. 2. Convergence phase where Rab5 positive domains are formed and converge towards one main domain. This phase is dependent on Rab7a positive endosome interaction that provides/supplies Rab7a to the maturing endosome. Finally, the converged domains leave the matured endosome to generate new Rab5 fully functional positive endosomes that will be primed for the next round of endosomal sorting through maturation (see text for detailed description).

## Materials and Methods

### Cell culture

Madine-Darby canine kidney strain II (MDCK) and HeLa cells were grown in an incubator at 37°C and 5 % CO_2_ in complete DMEM medium (Bio Whittaker), supplemented with 10% fetal calf serum (FCS, Integro), 2 mM L-Glutamine and 25 U/ml penicillin (all from Bio Whittaker). MDCK cells, stably transfected with invariant chain (Ii) in the pMep4 plasmid, where grown in the same conditions, with additionally 25 µg/ml Hygromycin (Bio Whittaker).

### DNA constructs

The cDNA encoding Ii (CD74) has been previously described (Bakke and Dobberstein, 1990). The subcloning of the Ii fragment into the pMep4 plasmid has been described (Gregers et al., 2003) and subsequently used (Skjeldal et al., 2012). The plasmid encoding mCherry-Rab5 (mCh-Rab5) has been previously described (Haugen et al., 2017). EGFP-Rab5Q79L was generously provided by Mitsonuri Fukuda (Tohoku University, Miyagi, Japan). pEGFP-Rab7a, pEGFP-Rab7aQ69L and pEGFP-Rab7aT22N were acquired from Cecilia Bucci’s laboratory (University of Salento, Lecce, Italy) and have been described previously (Bucci et al., 2000). The plasmid encoding dsRed-Rab7aT22N was a gift from Richard Pagano (Mayo Clinic and Foundation, Rochester, USA), through Addgene (plasmid #12662). The plasmid encoding mApple-Rab7a (Addgene plasmid #54945) was a gift from Michael Davidson (Florida State University).

### Cell Transfection

MDCK or HeLa cells were seeded in 6-well plates or in 35 mm glass bottom Petri dishes (MatTek Corporation). Transfections were performed overnight using Lipofectamine_®_ 2000 (Invitrogen), according to the manufacturer’s instructions. For each transfection, up to 1 µg DNA was used. The cells were imaged the day after the transfection.

### Expression of Ii (MDCK) and uptake of EGF (HeLa)

MDCK cells stably transfected with Ii-pMep4 were grown in 35 mm glass bottom Petri dishes (MatTek), in the presence of complete DMEM. To induce the expression of Ii, the cells were incubated with 25 µM CdCl_2_ overnight. Before imaging, the cells were washed with PBS. EGF-Alexa-Fluor-647 (Molecular Probes) was added to the HeLa cells, on the microscope, at a final concentration of 40 ng/ml.

### RNA interference

RNAi interference was used to knockdown Rab7a in Hela cells. Cells were plated one day before transfection in complete DMEM. Transfection was done with Oligofectamine_TM_ (Invitrogen), with 90nM siRNA targeting Rab7a or a non-targeting sequence for negative control. The following oligonucleotide sequences were used: Rab7a siRNA, sense sequence 5’-GGAUGACCUCUAGGUCAUCC-3’ anti sense sequence 5’-UUCUUCCUAGAGGUCAUCC - 3’(Spinosa et al., 2008)

### Western blotting and antibodies

Western blotting was used to determine the efficiency of the Rab7a knockdown. The silenced and control cells were lysed in a buffer containing 25 mmol Hepes, 125 mmol potassium acetate, 2,5 mmol magnesium acetate, and 5 mmol EGTA. Right before lysis, the buffer was complemented with 1 mM DTT, 0,5% NP-40 and a protease inhibitor cocktail (Roche). 20 µg of protein were loaded on precast Tris-Hepes gels (NuSep 12%) and transferred to Immobilon_®_ - P PVDF membranes (Millipore). The PVDF membranes were blotted with specific antibodies dissolved in a 2% milk solution, overnight at 4°C. The bands were visualized after incubation with ECL chemiluminescence kit (Amersham, GE Healthcare). An anti-tubulin antibody was used as a loading control (monoclonal mouse anti-a-Tubuline, Zymed, Thermofisher). The primary Rab7a antibody was purchased from Cell Signaling Technology (#2094S), and used 1:500. HRP-linked donkey anti-rabbit (NA934 GE Healthcare LifeSciences) were used as secondary antibody. Relative protein levels were measure with ImageJ.

## Live Confocal Microscopy

### Live cell Imaging and FRAP experiments

Live cell experiments were performed on four different microscopes; 1: Olympus iX81 FluoView 1000 inverted confocal microscope (Olympus, Hamburg, DE), equipped with a PlanApo 60X/1.30 oil objective, 2: Olympus SpinSR10 spinning disk confocal super resolution microscope equipped with a Yokogawa CSU-W1 SoRa. Images are acquired with a 60X Plan Apo 1.42 NA objective. 3: Zeiss LSM 880 equipped with and Airyscan detector and image acquisition with a 63X/1.40NA oil immersion objective. 4: The FRAP experiments were executed on an Andor Dragonfly equipped with a Mosaic FRAP module. The Andor Dragonfly is built on a Nikon TiE inverted microscope equipped a 60x/1.40 NA oil immersion objective. Bleaching of the EGFP-Rab5 was done with 405nm laser. Bleaching experiments were done at 100% laser power, 100ms bleaching time for the full-size endosome and 20ms for the converged Rab5 domains.

Obtained data from FRAP was normalized and corrected for bleaching (Pelkmans et al., 2001) and fitted by nonlinear regression to a function that assumes a single diffusion coefficient (Yguerabide et al., 1982);

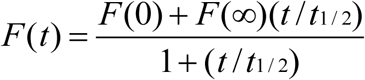

The values for *F(0), F(∞)* and *T*_*1/2*_ were calculated using GraphPad Prism 8 (http://www.graphpad.com/scientific-software/prism/) and the immobile fractions (IF) were calculated as described in Lippincott-Schwartz et *al (Lippincott-Schwartz et al*., *2001)*.

### Single analysis and quantification of D_1/2_

Single Rab5 and Rab7a positive endosomes in MDCK-Ii and Hela cells were manually tracked in ImageJ and mean intensity was measured over time. The result from 12-16 single endosomes per experiment was averaged over time with the variation depicted as SD in the graphs. D_1/2_ detachment was calculated performing a non-linear regression fit (Prism 8).

## Acknowledgements

We acknowledge the use of the NorMIC Oslo imaging platform at IBV. This work was supported by grants from the Norwegian Cancer Society and the Norwegian Research Council.

